# Within- and transgenerational plasticity of a temperate salmonid in response to thermal acclimation and acute temperature stress

**DOI:** 10.1101/2021.03.22.436503

**Authors:** Chantelle M. Penney, Joshua K. R. Tabh, Chris C. Wilson, Gary Burness

**Affiliations:** Environmental and Life Sciences Graduate Program, Trent University, Peterborough, Ontario, Canada, K9L 0G2; Ontario Ministry of Natural Resources and Forestry, Peterborough, Ontario, Canada, K9L 7B8; Department of Biology, Trent University, Peterborough, Ontario, Canada, K9L 1Z8

**Keywords:** Climate change, brook trout, metabolic rate, thermal tolerance, transgenerational acclimation, plasticity

## Abstract

Environmental temperatures associated with climate change are rising too rapidly for many species to adapt, threatening the persistence of taxa with limited capacities for thermal acclimation. We investigated the capacity for within- and transgenerational responses to increasing environmental temperatures in brook trout (*Salvelinus fontinalis*), a cold-adapted salmonid. Adult fish were acclimated to temperatures within (10□) and above (21□) their thermal optimum for six months before spawning, then mated in a full factorial breeding design to produce offspring from cold- and warm-acclimated parents as well as bidirectional crosses between parents from both temperature treatments. Offspring families were subdivided and reared at two acclimation temperatures (15□ and 19□) representing their current environment and a projected climate change scenario. Offspring thermal physiology was measured as the rate of oxygen consumption (MO_2_) during an acute change in temperature (+2□ h^-1^) to observe their MO_2_-temperature relationship. We also recorded resting MO_2_, the highest achieved (peak) MO_2_, and critical thermal maximum (CTM) as performance metrics. Within-generation plasticity was greater than transgenerational plasticity, with offspring acclimation temperature having demonstrable effects on peak MO_2_ and CTM. Transgenerational plasticity was evident as an elevated resting MO_2_ and the MO_2_-temperature relationship in offspring from warm-acclimated parents. Both parents contributed to offspring thermal responses, although the paternal effect was stronger. Although brook trout exhibit both within- and transgenerational plasticity for thermal physiology, it is unlikely that these will be sufficient for coping with long-term changes to environmental temperatures resulting from climate change.

**Summary:** Brook trout (*Salvelinus fontinalis*) exhibit within-generation and transgenerational plasticity for thermal performance, although neither response appears sufficient to cope with long-term climate change effects.

## INTRODUCTION

Environmental warming due to climate change is adversely affecting the physiology and persistence of species and populations globally (Moritz & Agudo, 2013; Whitney *et al*., 2016). With rising temperatures, thermal acclimation (within-generation plasticity, or WGP) allows for some short-term buffering of temperature effects through physiological adjustment but surviving climate change will likely depend on a population’s ability to move to more suitable habitats or adapt to warmer conditions (Habary *et al*., 2017). Species or populations that cannot migrate, or are restricted to localized habitats, are particularly vulnerable because they would be left to face temperatures higher than those to which they are physiologically capable of withstanding long-term (Somero, 2010). Although evolutionary change may provide the best option for long-term persistence (Munday *et al*., 2013), the accelerated rate of climate change is likely too rapid for most organisms to respond (Comte & Olden, 2017). This is especially true for those with long generation times or limited standing genetic variation (Willi *et al*., 2006; Munday *et al*., 2013; Meier *et al*., 2014).

One mechanism by which populations may compensate for long-term environmental change is via transgenerational plasticity (TGP; Jablonka *et al*., 1992; Bonduriansky *et al*., 2012; Norouzitallab *et al*., 2019) or epigenetic inheritance. TGP occurs when the inheritance of non-genetic factors and parental effects influence the offspring’s phenotype and physiological response to their environment (Deans & Maggert, 2015; Hanson & Skinner, 2016; Charlesworth *et al*., 2017). Environmental changes can likewise induce epigenetic changes (Deans & Maggert, 2015; Hanson & Skinner, 2016; Charlesworth *et al*., 2017) that can also be transmitted to the next generation and influence offspring phenotypes and physiology (Guillaume *et al*., 2016; Munday *et al*., 2017). By pre-conditioning their offspring for future environmental conditions, parents may buffer the impacts of environmental stressors and grant more time for the evolution of adaptive responses to occur (Bernatchez, 2016; Smith *et al*., 2016). Because TGP may provide populations with the ability to cope with chronic warming, TGP may represent an adaptive response to climate change (Greenspoon & Spencer, 2018; Yin *et al*., 2019; but see Sánchez□ Tójar *et al*., 2020).

Observations of TGP have been documented in a growing number of organisms (reviewed by Donelson *et al*., 2017; Yin *et al*., 2019). In fruit flies (*Drosophila melanogaster*), for example, thermal performance measured as climbing speed improved in offspring from parents reared in thermally variable conditions (Cavieres *et al*., 2019). Developmental rate increased in the marine polychaete (*Ophryotrocha labronica*) following multi-generational exposure to warming (Gibbin *et al*., 2017). In fish at warm ambient temperature, offspring from parents that had been acclimated to warm temperatures had a lower standard metabolic rate or increased maximum metabolic rate compared with offspring from cold acclimated parents (Donelson *et al*., 2012; Shama *et al*., 2014; Donelson *et al*., 2017). To date, most studies of TGP of aquatic vertebrates have focused on temperate or tropical species (Donelson *et al*., 2012; Salinas *et al*., 2012; Shama *et al*., 2014), however, high-latitude species are predicted to be most negatively impacted by climatic warming (Beitinger & Bennett, 2000).

Plasticity is thought to evolve in response to environmental variation (Somero, 2010, Colicchio & Herman, 2020), however, the extent to which WGP and TGP operate or interact in organisms that experience variable habitats remains unclear (Leimar & McNamara, 2015). For example, lake trout (*Salvelinus namaycush*) inhabit a thermally stable environment (Martin & Olver, 1980; Wilson & Mandrak, 2004) and exhibit little variation for thermal acclimation (WGP) across populations (McDermid et al., 2013; Kelly et al., 2014). In one of the few studies to investigate TGP in a cold-water species, Penney *et al*. (in press) showed that TGP was also limited in lake trout. In contrast to lake trout, brook trout (*S. fontinalis*) occupy different thermal habitats at different life stages (Biro *et al*., 2008; Smith & Ridgway, 2019), experience relatively high levels of environmental variation (Smith *et al*., 2020) and exhibit within generation thermal acclimation (McCormick *et al*., 1972; Stitt *et al*., 2014; Morrison *et al*., 2020). It is not yet known whether organisms like brook trout that display some variation in WGP for thermal tolerance are more or less capable of TGP.

The brook trout is a cold-adapted salmonid native to eastern North America found in cold (10-16□), well-oxygenated, freshwater habitats such as streams and lakes (Power, 1980; Smith & Ridgway, 2019). Brook trout also have a poor tolerance for warm temperatures (Beitinger & Bennett, 2000) making them highly vulnerable to climate change as temperatures become warmer and suitable habitat is lost (McKenna Jr, 2019). Thermal refugia in lakes are also being reduced as epilimnetic temperatures rise and the metalimnion shrinks (King *et al*., 1999); in some smaller lakes brook trout populations already encounter temperatures that push them to their physiological limits (21-23□; Smith *et al*., 2020) or prevent reproduction (Warren *et al*., 2012). The vulnerability of salmonids, including brook trout, to increases in temperature associated with climate change has resulted in a wealth of research on this taxon, yet their adaptive capacity, including transgenerational responses, to climate change remains largely unexplored.

In this study, we acclimated adult brook trout and their offspring to elevated temperatures to examine within-generation and transgenerational plasticity responses of offspring to a warming environment. We hypothesized that brook trout are capable of WGP and TGP responses to environmental temperatures, and that TGP would enhance upper thermal tolerance. To date, parental contributions to TGP have largely focused on the maternal environment which is thought to have a larger effect on offspring than the paternal environment (Shama *et al*., 2014; Best *et al*., 2018); however, paternal contributions are increasingly being reported across taxa (Hellman *et al*., 2020; Rutkowska *et al*., 2020). Our experimental design provided us with the opportunity to assess both maternal and paternal contributions to offspring thermal responses.

We measured offspring thermal physiology as the rate of oxygen consumption (MO_2_) during an acute change in temperature (+2□ h^-1^), to observe their MO_2_-temperature relationship. We also recorded resting MO_2_, the highest achieved (peak) MO_2_, and critical thermal maximum (CTM) as performance metrics. We first predicted that with acclimation to a warmer temperature, offspring peak MO_2_ and CTM would increase (demonstrating WGP; Morrison *et al*., 2020; Mackey *et al*., 2020), and offspring resting MO_2_ will be elevated due to the energetic cost of maintenance at a warmer temperature (Hartman & Cox, 2008; Norin & Metcalfe, 2019). Second, we predicted that offspring with warm-acclimated parents will have a lower resting MO_2,_ a higher peak MO_2_, and a higher CTM compared with offspring with cold-acclimated parents, because TGP would improve thermal tolerance at elevated temperatures (Donelson *et al*., 2012). Alternatively, resting MO_2_ could be higher in offspring from warm-acclimated parents, following the results of Penney *et al*. (in press). Lastly, we predicted that offspring and parental warm acclimation will interact (i.e., parents anticipate their offspring’s environment) to strengthen the effect of TGP on CTM, resting and peak MO_2_, and the MO_2_-temperature relationship.

## METHODS

All experiments were approved by the Trent University Animal Care Committee (Protocol # 24794) and the Ontario Ministry of Natural Resources and Forestry (OMNRF) Aquatic Animal Care Committee (Protocol FACC 136) and conducted according to the guidelines outlined by the Canadian Council on Animal Care.

The brook trout used for this study originated from wild spawn collections from a native population in Dickson Lake, Algonquin Provincial Park in south-central Ontario, Canada (45°47’ N, 78°12’ W), and have been kept in the OMNRF hatchery system since 2002 (OMNRF Fish Culture Stocks Catalogue 2005).

### Experimental design: Adult trout acclimation and breeding

Adult brook trout (age 5; 0.3-0.9 kg) from the Dickson Lake hatchery broodstock were transported to the OMNRF White Lake Fish Culture Station (Sharbot Lake, Ontario, Canada) in the spring of 2015 and implanted with PIT tags. A small caudal finclip (<0.25 cm^2^) was taken from each individual and separately stored in 95% ethanol to enable subsequent genetic parentage analysis of offspring families (described in supplemental material). In May 2015, adults were divided into two groups (n = 8 and 9, mixed sex), acclimated to one of two temperatures (10 ± 0.5 and 21 ± 0.5°C, respectively), and held until fall reproduction. The lower temperature was based on the temperature requirements for brook trout spawning, while the warmer temperature was selected to induce thermal stress without exceeding their physiological limits or compromising reproductive success (Hokanson *et al*., 1973; Blanchfield & Ridgway, 1997). Each group was kept in a 6000L flow-through tank covered with opaque acrylic lids to reduce stress to the fish, with light allowed in at the inflow and outflow to provide natural photoperiod cues. The tanks received water from White Lake (44°46’ N, 78°45’ W), and the desired temperatures were achieved by mixing inflows from above and below the lake’s thermocline. Fish were acclimated to these temperatures for the duration of the summer, after which the temperature of each tank followed the lake’s seasonal cooling from September to December. Tank water temperature and oxygen levels were checked daily, with temperature also logged every hour using two HOBO Tidbit loggers (Onset Computer Corporation, MA, USA) per tank for the duration of the adult acclimation period. The temperature data were collected from the loggers with a HOBO USB optic reader and HOBOware Pro (v. 2.3.0; Onset Computer Corporation, MA, USA) after spawning to track acclimation temperatures throughout the duration of the experiment.

Beginning in early October, the reproductive status of the trout was checked weekly by visual inspection following mild anesthesia (0.1 g L^-1^ MS-222; Aqua Life, Syndel Laboratories Ltd., B.C., Canada), and all adults were reproductive by December. As males and females came into reproductive condition, fish were dry-spawned by collecting gametes from anaesthetized fish, subdividing eggs from individual ripe females into two glass jars and fertilizing them with milt from separate males in a factorial cross mating design. In total, we used two males and two females from each of the two temperature treatments (8 adult fish in total) in two 2 x 2 factorial crosses, resulting in 16 families where the offspring were from parents of matched or mismatched thermal histories: C_♀_xC_♂_, C_♀_xW_♂_, W_♀_xC_♂_ and W_♀_xW_♂_ where C = cold and W = warm. Egg numbers for all families were equalized so that 140 mL of eggs from each female were sired by each male within each 2 x 2 factorial cross. Fertilized egg families were transported in insulated jars packed inside a cooler with ice packs to the OMNRF Codrington Fish Research Facility (Codrington, Ontario, Canada) where they were transferred to Heath trays receiving freshwater at ambient temperature (5-6□) under constant dim light for development.

### Experimental design: Offspring temperature acclimation

When fry reached the exogenous feeding stage, we randomly chose 20 offspring from each of the 16 families and divided them into two groups for acclimation to two different temperatures (15 and 19□). We chose the cooler acclimation temperature based on the optimal growth temperature reported for brook trout (McCormick *et al*., 1972), whereas the warm temperature simulated the potential warming due to climate change in the Great Lakes region by the end of the century (Hayhoe *et al*., 2010). Each group of 10 was moved into one of four larger (200 □) tanks: two tanks were designated for 15°C and the other two for 19°C so that each family had 10 representatives acclimated to each temperature. Each tank was separated into four sections to keep the families separate, but due to space constraints two families were kept in each tank section where the families sharing a section had a father in common. Individuals were identified to family after measurement trials by microsatellite genotyping (described in supplementary material).

Temperature acclimation began after the offspring were transferred to the larger tanks. We increased the water temperature at a rate of 1□ per day using titanium heaters (500W, model TH-0500, Finnex, IL, USA) with digital temperature controllers (model 192 HC 810M, Finnex, IL, USA) until the water in each tank reached its designated temperature (15 or 19°C). The temperatures were checked and recorded twice daily. During this time the fish were fed 5-6 times a day at 2-3% their body weight. The experiments began after the fish had been acclimated for at least 3-4 weeks.

### Respirometry set up

We explored the influence of parental thermal history on the MO_2_-temperature relationship in the offspring, the resting metabolic rate, peak metabolic rate and upper thermal tolerance of the offspring. The MO_2_-temperature relationship was measured as the metabolic rate of the offspring during an acute temperature increase of 2□ per hour, where metabolic rate was measured as the rate of oxygen consumption (MO_2_) with closed respirometry. Using this MO_2_ dataset, we determined the resting and peak metabolic rate of each individual offspring. Resting metabolic rate was recorded as the MO_2_ at the individual’s acclimation temperature before temperature increased with the acute temperature challenge (Chabot *et al*., 2016), and the peak MO_2_ as the individual’s highest MO_2_ during the trial. Finally, each individual’s upper thermal tolerance was measured as the critical thermal maximum (CTM) and was recorded as the temperature at which the fish could no longer maintain equilibrium following acute temperature increase.

Each experimental trial used eight custom built respirometers. Respirometers were made from a 8 cm diameter glass tube that was cut at a 4.5 cm length and sealed at one end (i.e. the floor of the chamber). The respirometer lids were made of acrylic. Each lid contained a fitting in the center for an O_2_ probe, and valves on opposite sides of the probe fitting to allow water to circulate through the respirometer chamber. Two respirometers were placed in each of four transparent plastic tubs and the tubs were seated on top of two, side-by-side stir plates (one plate per respirometer). The plates were used to spin a magnetic stir bar in each respirometer at approximately 60 RPM to prevent the establishment O_2_ gradients in the chambers and to keep water moving past an O_2_ probe (model DO-BTA, Vernier Software and Technology, OR, USA) that was inserted into the lid of each respirometer. The O_2_ probes were connected to a Lab Pro interface (Vernier Software and Technology) and O_2_ concentration within the respirometers was recorded using LoggerPro software (version 3.8.6; Vernier Software and Technology). Each respirometer also contained a perforated steel grid to separate the fish from the stir bar. Water from the tub was circulated through the respirometer at 4.5 L per minute using a submersible pump (universal type 1005, EHEIM GmbH & Co., Deizisau, Germany), and the water in each tub was also circulated with aerated, temperature-controlled freshwater from a source tank.

### Respirometry and critical thermal maximum protocol

Respirometry trials were conducted from August 9 to September 15, 2016. The night before a trial, eight fish were individually transferred into respirometers where they received a continuous flow of fresh water maintained at their acclimation temperature. They were left to adjust to the experimental apparatus overnight, and a thin sheet of black plastic covered each tub to minimize visual disturbance to the fish during the adjustment period and experimental trial. Fish were fasted for at least 12 hours prior to each trial to eliminate the physiological effects of digestion on the experimental results (Millidine *et al*., 2009).

We began measuring MO_2_ in each individual fish the next morning at 7:00. To measure MO_2_, the respirometer chambers were sealed by manually closing the respirometer valves and switching off the pumps that circulated water through the chambers. After a 30 second wait period, the reduction in chamber O_2_ concentration was recorded for 10 minutes. Afterwards, the flow valves were reopened to restore water circulation. Water temperature was then increased at a rate of 2□ per hour and the MO_2_ of each fish measured at each 1□ increase with 30 minutes between the repeated MO_2_ measurements. The rate of oxygen consumption (MO_2_) was calculated as,

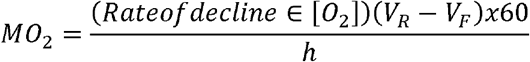

where *(Rate of decline in [O*_*2*_]) is the decline in water oxygen concentration (mg O_2_ L^-1^ min^-1^) during the 10-minute measurement period, *V*_*R*_ is the volume (L) of the respirometers, *V*_*F*_ is the volume of the fish (L) and *h* is the time in hours. The rate of decline was determined with LoggerPro, and we measured the linear fit of the drop in respirometer O_2_ concentration over time. If the linear correlation coefficient (r) was below 0.8, the datapoint was excluded from the analysis. This resulted in the exclusion of 225 out of 2,845 total datapoints collected. Some of these excluded values were measures of resting and peak MO2 (of 230 individuals, 43 resting MO_2_ and 5 peak MO_2_ were not included in the analysis).

The critical thermal maximum (CTM) for each fish was recorded as the temperature when it lost its righting response (i.e. equilibrium) and this was recognized as the point at which the fish could no longer maintain an upright position within the respirometer. All fish were closely monitored as temperature increased, and when a fish lost equilibrium it was quickly removed from the respirometer and euthanized with 0.3 g L^-1^ of tricaine methanesulfonate (MS-222; Aqua Life, Syndel Laboratories Ltd., B.C., Canada). Euthanized fish were immediately blotted dry on paper towels, measured for mass to the nearest 0.1 g and fork length (mm) using digital balance and calipers, respectively. Measurements of mass and length were used to calculate condition factor using the following formula,

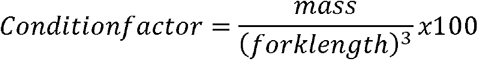

A tissue sample (caudal finclip) was taken from the euthanized fish and individually stored in 95% ethanol for microsatellite genotyping to identify each offspring to their respective family (described in supplementary material). Twelve of the 230 fish used for this experiment died at or just prior to collecting CTM, so were not included in the analysis of CTM.

To ensure that O_2_ would not become limiting at warmer temperatures, we monitored O_2_ saturation of the water throughout each trial. The source tank O_2_ concentration was kept at 6.0-7.0 mg L^-1^ and continuously checked with a YSI Pro probe (Hoskin Scientific, ON, Canada). If saturation levels lowered at high temperatures, O_2_ was supplemented to the source tank water using airstones and a tank of compressed O_2_ while ensuring that hyperoxia did not occur. Because the fish consumed O_2_ with increased rates at higher temperatures, we shortened the measurement period (<10 minutes) as necessary to avoid the O_2_ concentration from reaching the critical limit of 3.5 mg O_2_ L^-1^ during the MO_2_ measurement to avoid inducing a hypoxia response in the fish (Graham, 1949; Doudoroff & Shumway, 1970).

### Calculations and statistical analysis

The MO_2_ measured at the fish’s acclimation temperature before temperature began to rise with the acute temperature challenge was considered as the fish’s resting MO_2_. We report peak MO_2_ as the highest MO_2_ achieved during the respirometry trial. We do not report aerobic scope here because our measurement of peak MO_2_ may not necessarily represent the absolute maximum MO_2_ achievable by the offspring; max. MO_2_ is typically obtained using exhaustive exercise protocols which we did not use in this study. We analyze whole animal rates of O_2_ consumption with mass as a covariate rather than perform the analysis on mass-specific values because the former is statistically more appropriate (Hayes & Shonkwiler, 1996). The mean values reported from these models are referred to as mass-adjusted MO_2_. The raw data plotted as mass-specific MO_2_ is included in the supplementary material (Fig. S1).

The effect of parent and offspring acclimation temperatures on mass and condition factor was assessed using a general linear mixed effects model (GLMM). The models for mass and condition factor included offspring acclimation temperature (*T*_*O*_: cold or warm) and parental acclimation temperature (both parents combined into a single parental group: C_♀_xC_♂_, C_♀_xW_♂_, W_♀_xC_♂_ or W_♀_xW_♂_) as fixed effect predictors. An interaction term between offspring and parent acclimation temperature was also included as a fixed effect predictor to determine if parental acclimation temperature had differential effects on offspring mass and condition depending on whether the offspring were acclimated to a cold or warm temperature. Degree days was included as a random intercept to account for the potential effects of age on mass and condition factor. Degree days were calculated for each fish as the cumulative temperature experienced above 0°C (Chezik *et al*., 2013; Cook *et al*., 2018) until the beginning of the experimental trial. A Tukey’s HSD post-hoc analysis was performed if the test determined a significant effect of fixed predictors on mass or condition to uncover where differences occurred among pairwise comparisons.

To identify factors contributing to variation in resting MO_2_, peak MO_2_ and CTM, we evaluated competing statistical models using an Akaike Information Criterion corrected for small sample size (AICc). The possible model terms included offspring acclimation temperature (*T*_*O*_), maternal acclimation temperature (*T*_*M*_) and paternal acclimation temperature (*T*_*P*_) as fixed effect predictors, with interactions between all factors. Models also included offspring *Mass* as a covariate because the warm-acclimated offspring grew heavier than cold-acclimated ones and because metabolic rate scales with mass. The effects of maternal ID (*ID*_*M*_) and paternal ID (*ID*_*P*_) were included as random intercepts to control for statistical non-independence of offspring relatedness as some were full- or half-siblings based on the 2×2 factorial mating design. From the model AIC values, we calculated the ΔAIC, evidence ratio (ER) and Akaike weight (Wi) for each model and considered the best models as those with a ΔAIC ≤ 2 (Burnham & Anderson, 2002). We used the calculated metrics to compare the models and identify common parameters among the models that explained variation in resting MO_2_, peak MO_2_ and CTM.

The relationships between MO_2_ and temperature was curvilinear and could not be fit with a simple polynomial function. Instead, we fitted the relationship using a general additive model (GAM) as described in Penney *et al*. (in press). The GAM was generated by capturing the shape of the MO_2_ and acute challenge temperature relationship using all individuals with a cubic regression spline (i.e. GAM), and using just 7 knots to avoid over-fitting. From the GAM, we extracted the predicted MO_2_ of each offspring at each challenge temperature to use as a predictor in a general linear mixed model (GLMM). This method provided a workaround for the limitation of GAMs in accommodating complex interaction terms by accounting for the variation in the observed MO_2_ due to acute challenge temperature, allowing us to test whether the remaining variation in MO_2_ (i.e., that not due to acute challenge temperature) can be explained by offspring and parental acclimation temperature, and their interactions. To account for heterogeneity of variance in MO_2_ across acute challenge temperature we weighted the model by acute challenge temperature and included a type I autoregressive correlation structure (ρ = 0.221) to correct for autocorrelation between MO_2_ measurements as they occur at adjacent time-points during the acute temperature challenge.

To detect within-generation plasticity and trans-generational plasticity in the metabolic response of the offspring to an acute temperature challenge, we used a GLMM to test the effect of offspring, maternal and paternal acclimation temperature on the MO_2_-temperature relationship. We constructed a global model with the fixed effect terms of acute challenge temperature (*T*_*a*_, continuous variable), offspring acclimation temperature (*T*_*O*_; cold and warm), maternal acclimation temperature (*T*_*M*_; cold and warm), and paternal acclimation temperature (*T*_*P*_; cold and warm). All possible 2-way, 3-way and 4-way interactions between these terms were also included as fixed terms. As with our previously described models, the GLMM included *Mass* as a continuous predictor and the random intercepts of maternal and paternal ID (*ID*_*M*_ and *ID*_*P*_). Offspring ID was also included as a random intercept because repeated measurements of MO_2_ were collected from each individual at every 1□ increase in temperature.

All statistical analyses were conducted in JMP 13 (v. 18.1) or R (v. 3.5.2) with the level of significance set to 0.05. Linearity, homogeneity of variance, sample independence and residual normality were confirmed visually, and with the Shapiro-Wilk W, Levene’s and Brown-Forsythe tests. The factors that contributed to variation in body mass and condition factor were investigated using JMP 13. Statistical analyses of the resting and peak MO_2_, CTM, and MO_2_ during the temperature challenge were conducted using R with the ‘MuMIn’ (Barton, 2019), ‘lme4’ (Bates *et al*., 2015), ‘nlme’ (Pinheiro *et al*., 2019) and ‘mgcv’ (Wood, 2011) packages. We discovered that one of the peak MO_2_ datapoints was five standard deviations below the mean, therefore, this datapoint was not included in the final analysis.

## RESULTS

### Mass and condition factor

Warm-acclimated offspring were heavier overall compared with the cold-acclimated offspring (19□ offspring: 3.31 ± 0.08 g; 15□ offspring: 2.73 ± 0.08 g, GLMM: F_1,24.60_ = 25.22, p < 0.01). Parental acclimation temperature had a significant effect on offspring mass (GLMM: F_3,52.71_ = 13.78, p < 0.01). Offspring from parental groups C_♀_xC_♂_ and W_♀_xC_♂_ (3.50 ± 0.11 vs. 3.22 ± 0.11 g) were significantly heavier (p < 0.05) than those from the C_♀_xW_♂_ and W_♀_xW_♂_ parental groups (2.66 ± 0.12 vs. 2.69 ± 0.11 g), indicating that offspring with cold-acclimated fathers (C_♀_xC_♂_ and W_♀_xC_♂_) were heavier than those with warm-acclimated fathers. No other parental group comparisons were significantly different. There was no interaction between offspring acclimation and parental acclimation group (GLMM: F_3,52.71_ = 0.34, p = 0.80).

Warm-acclimated offspring had higher condition factor than cold-acclimated offspring (1.0 ± 0.01 vs. 0.96 ± 0.01; GLMM: F_1,23.25_ = 27.16, p < 0.01) and the condition factor of offspring was significantly affected by parental acclimation temperature (GLMM: F_3,53.89_ =6.10, p < 0.01). Offspring from parents that were both cold-acclimated (C_♀_xC_♂_) were in significantly better condition than offspring from parents that were both warm-acclimated (W_♀_xW_♂_; 1.0 ± 0.01 vs. 0.95 ± 0.01, respectively; p < 0.05). No other parental groups differed significantly from each other. There was no interaction between offspring acclimation temperature and parental acclimation group (GLMM: F_3,53.89_ = 0.13, p = 0.94).

### Critical thermal maximum

Critical thermal maximum was influenced by offspring acclimation temperature (WGP) but not transgenerational (i.e. parental) acclimation. Offspring acclimation temperature (*T*_*O*_) along with maternal and paternal ID (random effects: *ID*_*M*_ and *ID*_*P*_) best explained the variation in offspring CTM (ΔAIC ≤ 2, Table 1), no other model was within a ΔAIC ≤ 2. The effect of offspring acclimation temperature resulted in an approximately 0.4°C higher average CTM in warm-acclimated offspring versus cold-acclimated offspring (1.0 ± 0.01 vs. 0.96 ± 0.01; Fig. 1A and B).

**Table 1:** Summary of the top models determined with AIC_C_ to explain variation in brook trout offspring (age: 5 months) resting rate of oxygen consumption (MO_2_), peak MO_2_ and critical thermal maximum (CTM) with transgenerational acclimation.

| Measure | Model | $\Delta AIC$ | ER | $W_i$ | $R^2$ | Model |
| --- | --- | --- | --- | --- | --- | --- |
| CTM | 1 | 0 | 1 | 1 | 0.51 | $T_O + ID_M + ID_P$ |
| Resting $MO_2$ | 1 | 0 | 1 | 0.39 | 0.19 | $Mass + T_M + ID_M + ID_P$ |
| | 2 | 1.10 | 1.73 | 0.22 | 0.21 | $Mass + T_O + T_M + ID_M + ID_P$ |
| | 3 | 1.26 | 1.88 | 0.21 | 0.22 | $Mass + T_O + T_M + (T_O \cdot T_M) + ID_M + ID_P$ |
| | 4 | 1.49 | 2.11 | 0.18 | 0.17 | $Mass + T_M + T_P + ID_M + ID_P$ |
| Peak $MO_2$ | 1 | 0 | 1 | 0.48 | 0.52 | $Mass + T_O + ID_M + ID_P$ |
| | 2 | 1.02 | 1.67 | 0.29 | 0.51 | $Mass + T_O + T_P + ID_M + ID_P$ |
| | 3 | 1.53 | 2.15 | 0.23 | 0.53 | $Mass + T_O + T_M + ID_M + ID_P$ |

Offspring were from parents acclimated to either a cold or warm temperature and were similarly acclimated to cold or warm temperature. $T_P$ , $T_M$ , $T_O$ are the paternal, maternal and offspring acclimation temperatures, respectively, and $ID_M$ and $ID_P$ are the maternal and paternal individual identification (random effects).

**Fig. 1:**
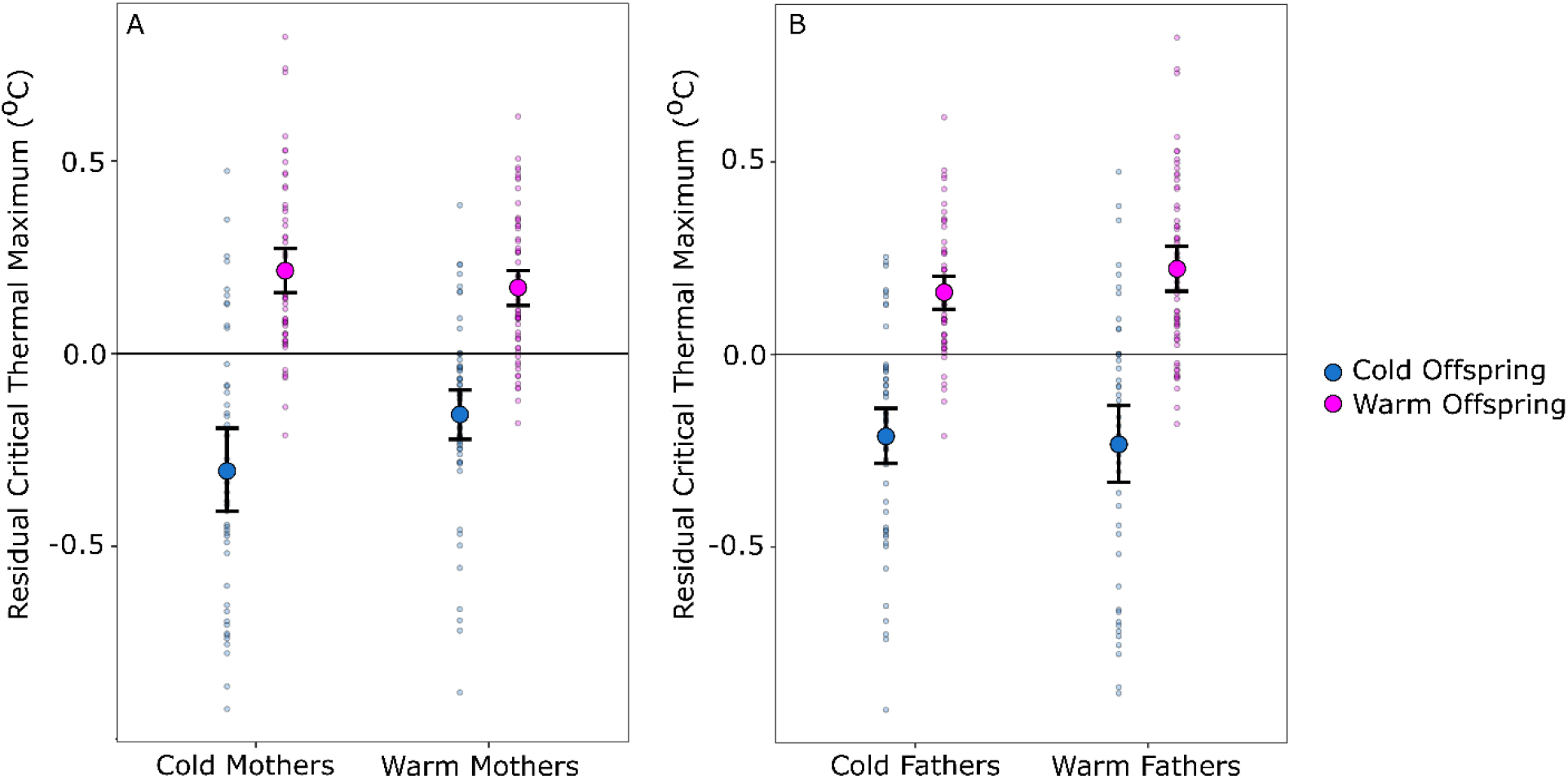
The effect maternal (A) and paternal (B) acclimation temperature on the critical thermal maximum (CTM) of brook trout offspring (age: 5 months) acclimated to a cold (15□, n = 100) or warm (19□, n = 116) temperature. Values represent the residuals (± confidence intervals) from the population mean, controlling for the effect of mass and family.

### Resting and peak metabolic rate

Resting MO_2_ was affected by offspring and parental acclimation temperature. Four models best explained variation in resting MO_2_, and each included *Mass* and maternal acclimation temperature (*T*_*M*_), with maternal and paternal ID (*ID*_*M*_ and *ID*_*P*_) as random effects (Table 1). The first model contained only these factors, while offspring acclimation temperature (*T*_*O*_) appeared in models 2 and 3 with an interaction occurring between offspring and maternal acclimation temperature in model 3 (Table 1). Paternal acclimation temperature (*T*_*P*_) appeared only once in the top four models and occurred in model 4, which was 2.11 (ER) less likely to best explain variation in the data when compared with model 1 (Table 1). Plotting the resting MO_2_ residuals showed that, in general, warm-acclimated offspring tended to have residual resting MO_2_ values slightly below the population mean (Fig. 2 A & B). With regard to parental acclimation temperature, the residual resting MO_2_ of cold-acclimated offspring from warm-acclimated mothers (Fig 2A) and warm-acclimated fathers (Fig 2B) was higher than the population mean. In contrast, when offspring from warm-acclimated mothers (Fig 2A) and warm-acclimated fathers (Fig 2B) were warm-acclimated, offspring resting MO_2_ was lower than the population mean. Together this suggests a transgenerational effect of lowering resting MO_2_ when parents and offspring each experience warming. The interaction between maternal and offspring acclimation temperature (*T*_*O*_ *· T*_*M*_) was evident in the residual plot (Fig. 2A): when mothers were cold-acclimated the residual resting MO_2_ of their cold-acclimated offspring was slightly lower than that of their warm-acclimated offspring, this trend reversed when the mothers were warm-acclimated.

**Fig. 2:**
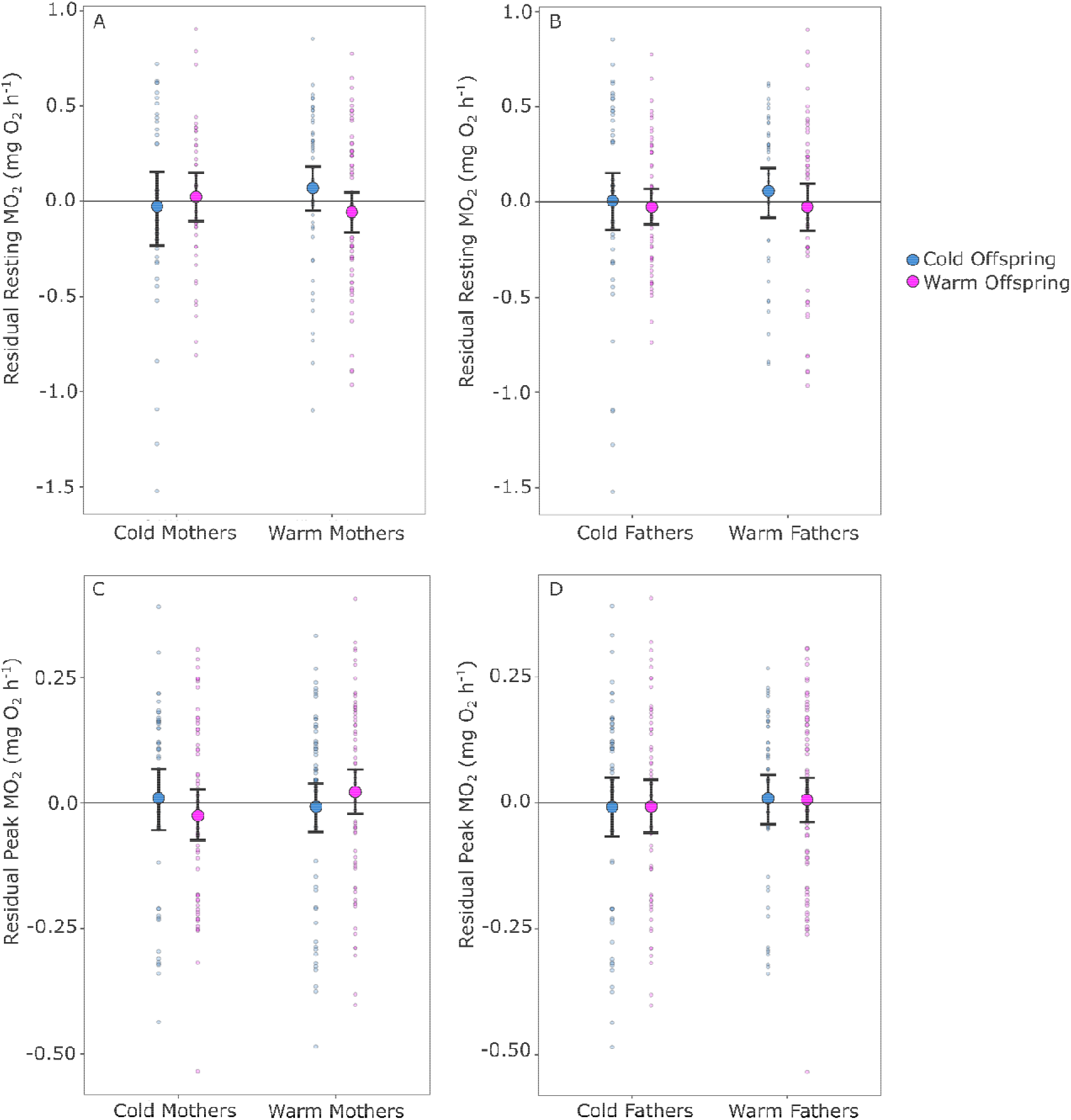
The effect of maternal (A and C) and paternal (B and D) acclimation temperature on the resting rate of oxygen consumption (MO_2_), and peak MO_2_ of brook trout offspring (age: 5 months) acclimated to a cold (15□) or warm (19□) temperature (n = 85-122). Values represent the residuals (± confidence intervals) from the population mean, controlling for the effect of mass and family.

Peak MO_2_ was also affected by offspring and parental acclimation temperature. Three models best explained variation in peak MO_2_ (ΔAIC ≤ 2), each with maternal and paternal ID (*ID*_*M*_ and *ID*_*P*_) as random effects (Table 1). *Mass* and offspring acclimation temperature (*T*_*O*_) were the best predictors of peak MO_2_ as these two fixed effects factors occurred in all three models and were the only fixed parameters in the first model. Paternal (*T*_*P*_, model 2) and maternal (*T*_*M*_, model 3) acclimation temperature each appeared only once among the three models. Plots of the residual peak MO_2_ showed only slight differences due to offspring acclimation temperature, being marginally lower in warm-acclimated offspring (Fig. 2C & D). Warm-acclimated mothers and warm-acclimated fathers slightly elevated the peak MO_2_ in their offspring (Fig. 2 C & D).

### Metabolic response of offspring to an acute temperature challenge

Offspring mass had a significant effect on oxygen consumption, as rates were higher in heavier fish (*Mass*: p < 0.001; Table 2). Offspring MO_2_ also increased with challenge temperature (*T*_*a*_: p < 0.001; Table 2) and the MO_2_-temperature relationship was similar between offspring of cold and warm acclimation temperatures (*T*_*O*_: p = 0.880; Table 2).

**Table 2:** Factors contributing to variation in brook trout rate of oxygen consumption (MO_2_).

| Parameter | Coefficient | S. E. | DF | t-value | p-value |
| --- | --- | --- | --- | --- | --- |
| *Intercept* | -0.84 | 0.11 | 17.87 | 7.48 | <0.001 |
| *Mass* | 0.20 | 0.02 | 182.14 | 12.27 | <0.001 |
| $T_a$ | 1.19 | 0.09 | 2338.40 | 13.03 | <0.001 |
| $T_O$ | 0.02 | 0.12 | 483.13 | 0.15 | 0.880 |
| $T_M$ | 0.22 | 0.11 | 180.89 | 2.05 | 0.042 |
| $T_P$ | 0.44 | 0.14 | 11.08 | 3.08 | 0.010 |
| $T_O \cdot T_M$ | 0.011 | 0.15 | 545.40 | 0.07 | 0.941 |
| $T_O \cdot T_P$ | -0.05 | 0.16 | 869.28 | 0.30 | 0.766 |
| $T_M \cdot T_P$ | -0.22 | 0.16 | 943.48 | 1.38 | 0.168 |
| $T_a \cdot T_O$ | -0.003 | 0.12 | 2339.95 | 0.02 | 0.981 |
| $T_a \cdot T_M$ | -0.17 | 0.12 | 2337.07 | 1.43 | 0.152 |
| $T_a \cdot T_P$ | -0.36 | 0.13 | 2338.77 | 2.70 | 0.007 |
| $T_O \cdot T_M \cdot T_P$ | -0.15 | 0.22 | 646.15 | 0.70 | 0.487 |
| $T_a \cdot T_O \cdot T_M$ | -0.13 | 0.17 | 2345.00 | 0.78 | 0.435 |
| $T_a \cdot T_O \cdot T_P$ | -0.05 | 0.18 | 2343.07 | 0.27 | 0.784 |
| $T_a \cdot T_M \cdot T_P$ | 0.17 | 0.18 | 2338.35 | 0.94 | 0.348 |
| $T_a \cdot T_O \cdot T_M \cdot T_P$ | 0.30 | 0.24 | 2344.12 | 1.27 | 0.205 |

Offspring (age: 5 months) from parents acclimated to either a cold or warm temperature were similarly acclimated to cold or warm temperature. $T_P$ , $T_M$ , $T_O$ are the paternal, maternal and offspring acclimation temperatures, respectively, and $T_a$ represents the acute temperature challenge (the cubic regression spline).

There was a transgenerational effect of parental acclimation temperature on offspring MO_2_ responses to an acute temperature challenge. The acclimation temperature of the mothers and fathers (Fig. 3A & B, respectively) each significantly affected the offspring’s metabolic response to the challenge temperature. That is, the MO_2_ of the offspring from warm-acclimated parents was elevated compared with offspring from cold-acclimated parents (Fig. 3A & B). While the effect was significant for both parents, a stronger effect occurred on the paternal side (*T*_*M*_: p = 0.042; *T*_*P*_: p = 0.010; Table 2).

**Fig. 3:**
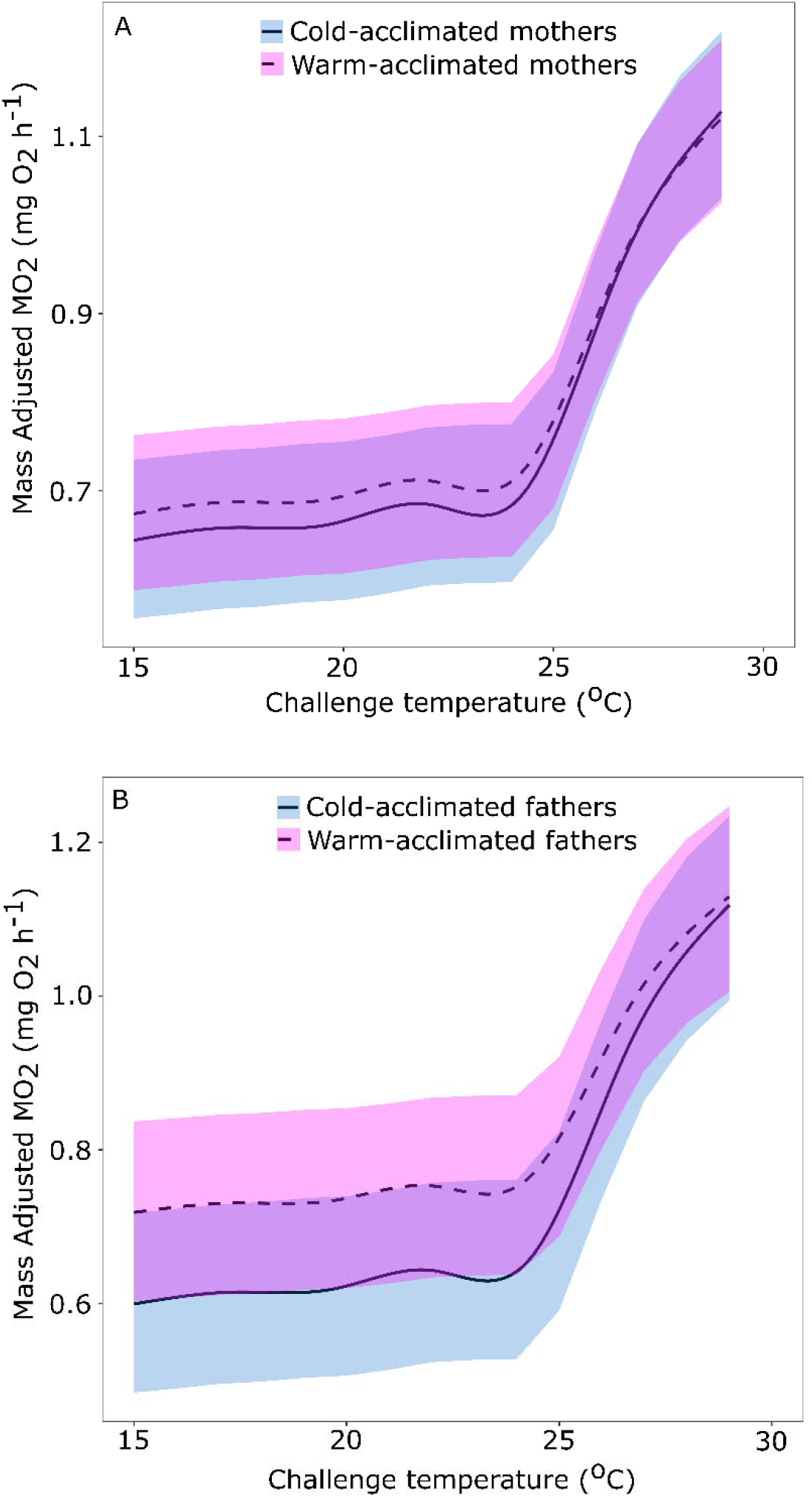
The influence of A) maternal and B) paternal acclimation temperature on the change in the rate of oxygen consumption (MO_2_) of cold-(15□, n = 105) and warm-(19□, n = 125) acclimated brook trout offspring (age: 5 months) given an acute temperature challenge of +2°C·h^-1^. Plots show means and 95% confidence intervals for cold- and warm-acclimated offspring shown in blue and red respectively as estimated from the GLMM where challenge temperature corresponds to a spline. Rates of oxygen consumption (MO_2_) were statistically-adjusted for effects of body mass (see text).

There was a significant statistical interaction between challenge temperature and paternal acclimation temperature (*T*_*a*_ *· T*_*P*_: p = 0.007; Table 2). Offspring mass-adjusted MO_2_ was lower in offspring from cold-acclimated fathers compared to those from warm-acclimated fathers when challenge temperatures were below 25□, but the groups tended to converge at temperatures above 25□ (Fig. 3B). No other interaction terms were significant.

## DISCUSSION

We saw evidence of WGP and TGP in brook trout upper thermal tolerance and metabolic rate with greater WGP and TGP effects. We found partial support for the prediction that CTM, resting MO_2_ and peak MO_2_ would all increase when offspring were acclimated to a warmer temperature. As predicted, acclimating offspring to a warmer temperature resulted in slightly (0.4□) higher mean CTM. In contrast, warm-acclimation of the offspring had a modest effect of lowering resting MO_2_, with offspring acclimation temperature appearing in half of the top AIC models for resting MO_2_. Similarly, peak MO_2_ was slightly lower in warm-acclimated offspring. Taken together, these results demonstrate within-generation thermal plasticity in the juvenile brook trout, however, plasticity was more limited than expected based on previous studies (e.g., McCormick *et al*., 1972; Stitt *et al*., 2014; Morrison *et al*., 2020). Limited within-generation plasticity was further supported by the results of the acute temperature challenge, where the MO_2_-temperature relationship did not differ between the two offspring acclimation temperatures (*T*_*O*_).

Our results partially supported the prediction that TGP would improve thermal tolerance at elevated temperatures (Table 1). Parental acclimation temperature had no effect on offspring CTM, but both maternal and paternal acclimation temperature influenced offspring resting MO_2_, and to a lesser extent peak MO_2_. Maternal and paternal acclimation temperature each contributed to the offspring’s response to the acute temperature challenge, with an overall upward shift in the curve for offspring from warm-acclimated parents. To our surprise, paternal TGP contribution to offspring thermal performance (MO_2_-temperature relationship) was larger than the maternal contribution.

### Within-generation plasticity

Thermal plasticity is thought to have evolved when organisms experience temperature fluctuations within their lifetimes (Beaman *et al*., 2016). We expected brook trout to show some within-generation plasticity for thermal tolerance based on their use of different habitats with varying thermal conditions throughout their life (Biro *et al*., 2008; Smith and Ridgway, 2019). Additionally, there is evidence that different populations have differing capacities for thermal acclimation (McDermid *et al*., 2012; Stitt *et al*., 2014).

Within-generation plasticity was observed in CTM; CTM increased with offspring acclimation temperature. CTM was in the expected range for brook trout acclimated to our temperatures (Wehrly *et al*., 2007; O’Donnell *et al*., 2020), however, plasticity in their upper thermal tolerance appears limited. The CTM only increased by ∼0.4□ in warm-acclimated offspring despite the 4□ difference in acclimation temperature between the two groups of offspring. Though the rate of heating used in our experiment (+2 □/hr) has been deemed appropriate rate for measuring CTM in brook trout (Galbreath *et al*., 2004), it is unclear whether the CTM of warm-acclimated offspring would be higher with a slower rate of heating. For example, Morrison *et al*. (2020) found that brook trout acclimated to 15□ and 20□ had significantly different CTM when heating the fish at a rate of 0.3□ per minute (i.e, +18 □/hr).

We detected an effect of thermal acclimation on offspring resting MO_2_, however, the effect was small (Table 1). Resting metabolic rate typically increases with acclimation temperatures until the individual reaches its *pejus* temperature (approximately 20□ for brook trout: Hartman & Cox, 2008). Thus, we anticipated that resting MO_2_ would be higher in warm-acclimated offspring compared to cold-acclimated offspring, largely because they were being measured at a warmer temperature, but we did not see this trend. It is not immediately clear why resting MO_2_ was not more elevated in warm-acclimated offspring compared with cold-acclimated offspring. A previous study on variation in upper thermal tolerance and metabolic rate of brook trout found that individuals originating from Dickson Lake (the same lake from which our brook trout originated) had a higher standard metabolic rate (SMR) following acclimation to 16□ compared to 20□, though the authors did not suggest the lower SMR at 20□ was due to the fish reaching their *pejus* temperature (Stitt *et al*., 2014). Stitt *et al*.’s (2014) experimental temperatures were comparable to ours (15-16□ and 19-20□), however, the fish tested were of different life stages (yearling vs. adult). The factors influencing resting MO_2_ (or SMR) of brook trout warrants further investigation.

Offspring acclimation temperature had a modest effect on peak MO_2_, demonstrating some within-generation plasticity in this parameter. Some fish species are capable of extending the upper limit of MO_2_ (i.e. peak MO_2_) when acclimated to warmer temperatures (reviewed by Schulte, 2015). For example, exercise-induced maximum metabolic rate increased by 20-30% in lake trout acclimated from 8 to 15□ (Kelly *et al*., 2014). It is important to note that our measurement of peak MO_2_ is not equivalent to maximum MO_2_ which is typically measured in the fish after exhaustive exercise or a chase protocol (as in Kelly *et al*., 2014). Our results suggest that brook trout peak MO_2_ is capped within the values we observed in our study. This generally agrees with the idea that metabolic ceilings, like peak MO_2_ or CTM, are relatively thermally (acclimation) insensitive (Sandblom *et al*., 2016; Norin & Metcalfe, 2019; Morrison *et al*., 2020).

Offspring acclimation temperature did not influence the offspring’s MO_2_ response to an acute temperature challenge (+2□ h^-1^). Interestingly, offspring MO_2_ did not begin to rise until the challenge temperature exceeded 23°C. Although the reason for the sudden increase in MO_2_ at ∼23□ is not immediately clear, it could be due to a physiological stress response(s) being initiated at this temperature, especially considering that 23□ is near the upper incipient lethal temperature recorded for these fish (24□; Fry *et al*., 1946; Wehrly *et al*., 2007). Identifying the physiological processes that result in the increased MO_2_ at ∼23□ in brook trout would require further study, although previous works have reported on correlated mechanisms during an acute thermal challenge. Sandblom & Axelsson (2007) demonstrated an increase in hematocrit, likely resulting from splenic contraction, in rainbow trout when challenged with an acute temperature increase. Splenic contraction would increase the number of red blood cells available to meet the increased oxygen delivery demands associated with acute warming (Clark *et al*., 2008). Chadwick *et al*. (2015) saw that levels of HSP70 and glucose increased in juvenile brook trout when challenge temperatures reached approximately 21□. It is possible that stress responses, such as induction of molecular chaperones or mobilization of energetic resources were initiated at 23□ in the juvenile brook trout in our study, thus increasing metabolic rate at this temperature.

### Transgenerational plasticity

Both maternal and paternal acclimation temperature affected the offspring’s MO_2_- temperature relationship, with an overall upward shift for offspring from warm-acclimated parents. This was also reflected in offspring resting MO_2_ and it implies that offspring will incur a higher cost of living (Norin & Metcalfe, 2019) when parents are warm-acclimated. It is thought that TGP is adaptive, thus selected for, when the environment changes predictably across generations (Jablonka *et al*., 1992; Bonduriansky *et al*., 2012; Norouzitallab *et al*., 2019; but see Uller *et al*., 2013). Based on this idea, TGP would be predicted to be weak in stenothermal organisms that have adapted to relatively cold, thermally stable habitats. The limited available evidence supports this: in lake trout (*S. namaycush*), a cold-adapted stenothermal congener of brook trout, TGP was limited, and most evident as elevated MO_2_ in warm-acclimated offspring from warm-acclimated parents (Penney *et al*. in press). In contrast, in eurythermal or warm-adapted fish species, metabolic rates are *reduced* in warm-acclimated offspring from warm-acclimated parents compared to cold-acclimated parents (Donelson *et al*., 2012; Shama *et al*., 2014; Donelson *et al*., 2017).

In our study, peak MO_2_ varied only slightly with maternal or paternal acclimation temperature, and no transgenerational effect on offspring CTM was detected. Our results agree with the limited number of studies on the transgenerational effect of warming on maximum metabolic rates and CTM. Sandblom *et al*. (2016) found that European perch (*Perca fluviatilis*) held at higher temperatures for multiple generations displayed little change in their maximum rate of O_2_ consumption and CTM. Similarly, the CTM of largemouth bass (*Micropterus salmoides*) did not differ after multi-generational exposure to warmer temperature (White & Wahl 2020). Further, CTM and peak MO_2_ of lake trout offspring were also not affected by parental exposure to a warmer environment (Penney *et al*., in press), which is perhaps unsurprising given that there was limited WGP in lake trout peak MO_2_ and CTM with offspring acclimation in the same study. Together, these studies suggest TGP is unlikely to significantly alter CTM or peak MO_2_ in response to increased environmental temperatures over relatively short multi-generation timespans, reinforcing evidence that these metabolic ceilings are likely to be exceeded in ecological timeframes (Sandblom *et al*., 2016; Norin & Metcalfe, 2019; Morrison *et al*., 2020).

We sought to examine the extent to which WGP and TGP operate or interact in brook trout. While WGP was evident in peak MO_2_ and CTM, it was through TGP that warm-acclimated parents elevated resting MO_2_ and affected the MO_2_-temperature relationship in offspring. The importance of TGP relative to WGP may depend on life stage and variation in the habitat experienced at each life stage. It is possible that transgenerational effects are strongest in early-juvenile life stages (Yin *et al*., 2019), but only when the environment is stable. In contrast, WGP may be the favoured strategy when temperatures are more variable (Leimar & McNamara, 2015). In fact, in situations where environmental temperature variation exists, TGP effects may be overridden by WGP, as found in stickleback (Shama, 2017). Our study examined juvenile brook trout 5-6 months after hatching, at a time when they would be feeding in shallow depths near shore and near the surface in warmer water (Biro *et al*., 2008). It would be interesting to examine the relative influences of WGP and TGP at earlier or subsequent life stages. We are not aware of any such studies to date, although experiments examining WGP and TGP at multiple life stages could be very informative.

### Relative parental contributions

Although both maternal and paternal thermal history (temperature acclimation) each contributed to offspring thermal physiology, we did not find strong evidence that transgenerational effects were additive (i.e., stronger when the offspring had a warm mother *and* a warm father). Maternal and paternal acclimation temperature (*T*_*M*_ + *T*_*P*_) appeared in the same model only once for resting MO_2_ (model 4, Table 1), but not for peak MO_2_ or CTM. Similarly, each parent contributed to their offspring’s MO_2_ response to an acute temperature increase.

Paternal effects have received less attention relative to maternal effects (Rutkowska *et al*., 2020) and the size of the epigenetic paternal contribution to such changes relative to the maternal contribution is still debated (reviewed by Best *et al*., 2018). In the few studies that have tested relative parental contributions to TGP in metabolic traits in fish, the paternal contribution is either less than (Shama *et al*., 2014) or comparable to the maternal contribution (Penney *et al*., in press). In this study, fathers surprisingly appeared to have greater contributions to TGP than did mothers. Paternal effects are complex, can depend on the sex of the offspring, and can vary depending on the environment experienced by paternal grandparents (Crean & Bonduriansky, 2014; Hellman *et al*., 2020). Environmentally-mediated epigenetic changes do occur in sperm (Ord *et al*., 2020; Immler, 2020) and these along with cytoplasmic components can influence offspring phenotypes (summarized by Donkin & Barrès, 2018). Parents can also have opposing effects on gene expression in their offspring despite both parents having received the same treatment, in that a gene may be maternally downregulated but paternally upregulated in the offspring (Bautista *et al*., 2020). While epigenetic regulation of gene expression may be an underlying factor in the paternal contribution we observed in our study, we are not aware of another study showing such a large paternally-mediated TGP contribution to thermal responses relative to the maternal contribution.

## Summary

The brook trout in our study displayed a limited transgenerational response to warmer temperature. Transgenerational effects on offspring phenotypes depend on genotype or ecotype (Verhoeven & van Gurp, 2012; Vayda *et al*., 2018), and TGP is predicted to arise in populations that experience variation in temperature over multiple generations (Beaman *et al*., 2016; Yin *et al*., 2019). Because different populations of brook trout display variation in thermal tolerance and capacity for acclimation (McDermid *et al*., 2012; Stitt *et al*., 2014) it would be informative to assess whether transgenerational responses to warming vary among stream and lake populations, and across the species’ range. One might predict, for example, that daily as well as seasonal thermal variation in stream environments (Chadwick & McCormick, 2017) would select for increased TGP compared with lake habitats. Like many other cold-adapted organisms, brook trout are being threatened by climate change-induced habitat impacts (King *et al*., 1999; Robinson *et al*, 2010; Bassar *et al*., 2017). Although previous studies have suggested TGP may buffer the impact of environmental stressors and facilitate an evolutionary response to climate change (Bonduriansky *et al*., 2012; Bernatchez, 2016; Smith *et al*., 2016), the limited response observed in our study suggests that TGP may not sufficiently enable brook trout to keep pace with anticipated changes in environmental temperatures.

## Acknowledgements

Adult brook trout were provided by the Ontario Ministry of Natural Resources and Forestry (OMNRF) Fish Culture Section, and fish care, rearing space and logistical support was provided by the OMNRF White Lake Fish Culture Station. Bill Sloan, Scott Ferguson, Anne McCarthy (OMNRF) and John Dewart assisted with breeding adult fish. Bill Sloan and Scott Ferguson kindly provided logistical support for all aspects of fish husbandry and juvenile fish care, and assisted with the experiments conducted at the OMNRF Codrington Fisheries Research Facility. Caleigh Smith (OMNRF) genotyped the fish for identification of juveniles to family. Kurt Gamperl (Memorial University) provided advice on respirometer design, and Ben Bolker and John Dushoff (McMaster University) advised on statistical analysis.

